# A Diffusion-Based Framework for Designing Molecules in Flexible Protein Pockets

**DOI:** 10.1101/2025.05.27.656443

**Authors:** Jian Wang, Dong Yan Zhang, Shreshty Budakoti, Nikolay V. Dokholyan

**Affiliations:** Department of Neuroscience and Experimental Therapeutics, Penn State College of Medicine, Hershey, PA, 17033-0850, USA; Department of Biomedical Engineering, Pennsylvania State University, University Park, PA; Department of Chemistry, Pennsylvania State University, University Park, PA

**Author notes:** Corresponding author: Nikolay V. Dokholyan,.

## Abstract

The design of molecules for flexible protein pockets represents a significant challenge in structure-based drug discovery, as proteins often undergo conformational changes upon ligand binding. While deep learning-based approaches have shown promise in molecular generation, they typically treat protein pockets as rigid structures, limiting their ability to capture the dynamic nature of protein-ligand interactions. Here, we introduce YuelDesign, a novel diffusion-based framework specifically developed to address this challenge. YuelDesign employs a new protein encoding scheme with a fully connected graph representation to encode protein pocket flexibility, a systematic denoising process that refines both atomic properties and coordinates, and a specialized bond reconstruction module tailored for de novo generated molecules. Our results demonstrate that YuelDesign generates molecules with favorable drug-likeness and low synthetic complexity. The generated molecules also exhibit diverse chemical functional groups, including some not even present in the training set. Redocking analysis reveals that the generated molecules exhibit docking energies comparable to native ligands. Additionally, a detailed analysis of the denoising process shows how the model systematically refines molecular structures through atom type transitions, bond dynamics, and conformational adjustments. Overall, YuelDesign presents a versatile framework for generating novel molecules tailored to flexible protein pockets, with promising implications for drug discovery applications.

## INTRODUCTION

The design of three-dimensional molecular structures with specific properties remains a fundamental challenge in drug discovery. In recent years, generative artificial intelligence techniques, especially autoregressive models^1^ and diffusion models^2^, has transformed *de novo* drug discovery^3^ by facilitating the efficient generation of novel molecular structures. These generative approaches are broadly divided into two main categories: 2D molecular topology generation, which involves constructing molecular graphs or string-based representations such as SMILES^4^ and SELFIES^5^, and 3D molecular conformation generation, which aims to predict atomic positions in three-dimensional space.

In the domain of 2D molecular generation, commonly employed approaches include autoregressive models^6–8^ (e.g., RNNs for SMILES/SELFIES generation), variational autoencoders^9^ (VAEs) (GraphVAE^10^ and CGVAE^11^), generative adversarial networks (GAN)-based models like MOLGAN^12^, and flow-based frameworks including GraphNVP^13^ and MoFlow^14^. Concurrently, 3D molecular generation has made notable progress through a range of innovative approaches. For example, DiffSBDD^15^ showcased the use of diffusion models in structure-based drug design by leveraging equivariant graph neural networks^16^ (EGNN) to generate candidate molecules within protein binding pockets. Other notable diffusion-based methods include DiffSMol^17^, DiffBP^18^, and TargetDiff^19^.

A critical yet underexplored challenge in structure-based molecular design is the treatment of protein binding pockets during the generation process. This challenge is analogous to the distinction between rigid and flexible docking^20,21^ in protein-ligand docking studies. While rigid docking simplifies the binding site as a static structure, flexible docking allows for conformational adjustments, enabling more accurate predictions of binding modes and affinities. This concept, known as induced fit^22,23^, underscores how ligand binding can trigger structural rearrangements within the protein pocket, potentially exposing or altering key interaction sites. Whether to account for induced fit also significantly impact downstream tasks such as binding affinity prediction^24,25^, virtual screening^26^, and target identification^27^. Despite its significance, the dynamic nature of protein pockets remains largely overlooked in current molecular design frameworks. Approaches like DiffSBDD typically treat the binding pocket as a rigid entity.

In this work, we present YuelDesign, a novel diffusion-based molecular design framework (Figure 1) specifically developed to address the challenge of designing molecules for flexible protein pockets. Our framework integrates several key innovations to tackle this complexity. First, we introduce a new protein pocket encoding scheme (Methods), in which side chain features are not explicitly encoded, with a fully connected graph representation to effectively account for pocket flexibility. Second, we implement a systematic denoising process that refines both atomic properties and molecular structures, facilitating the learning of chemical constraints throughout the generation process. Third, we incorporate a post-generation bond reconstruction module, YuelBond^28^, specifically designed to restore bond integrity in *de novo* generated molecules, ensuring chemical validity and structural stability.

**Figure 1.**
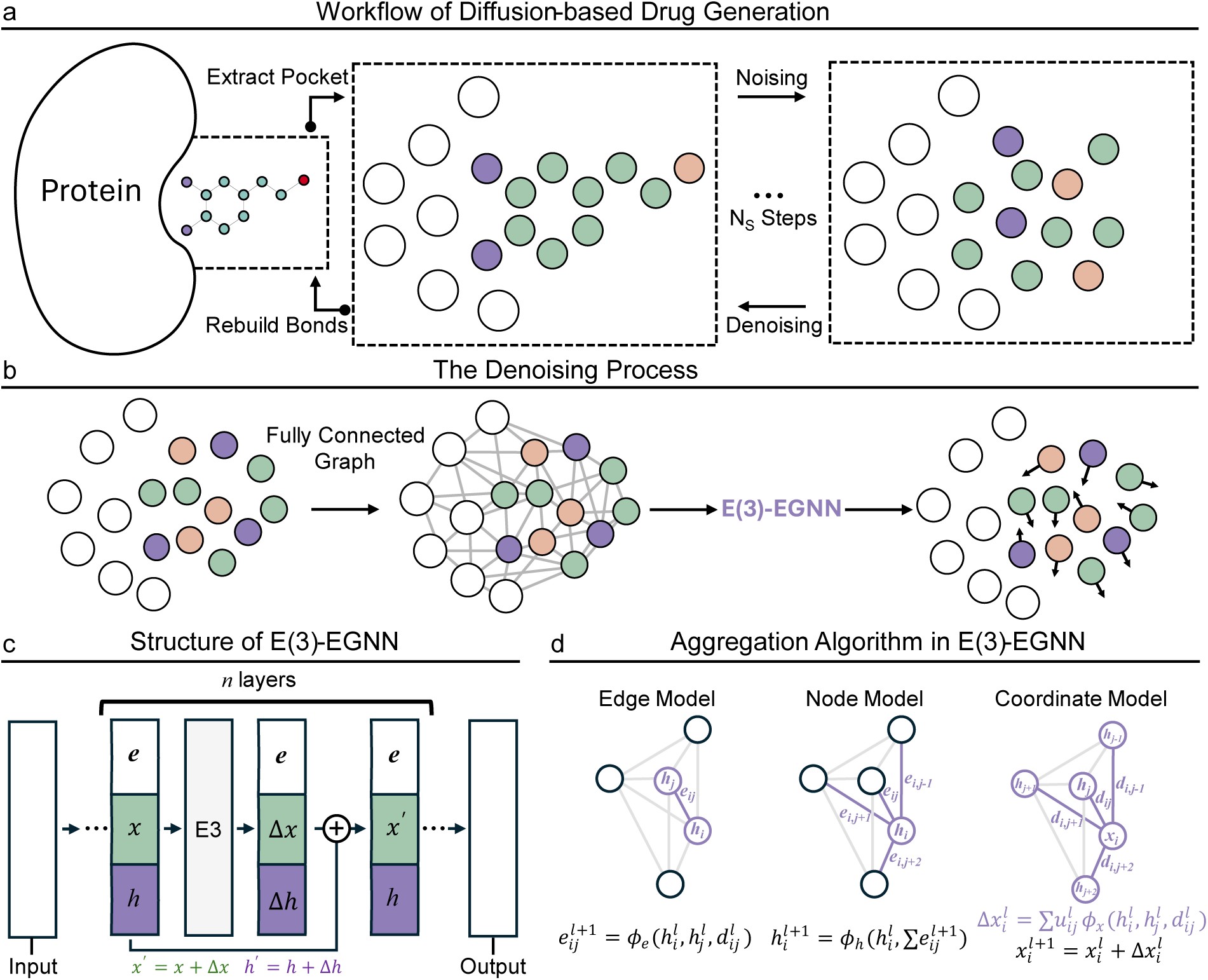
Workflow and architecture of YuelDesign. (a) The pipeline begins with protein pocket extraction, followed by iterative noising (Nₛ steps) and denoising to generate 3D molecular conformations. Bond reconstruction is applied post-denoising. (b) Denoising via an E(3)-equivariant Graph Neural Network (E(3)-EGNN). (c) The E(3)-EGNN architecture comprises *n* layers with dedicated edge, node, and coordinate models. (d) Message aggregation mechanism within E(3)-EGNN, updating atom positions (x) and features (h) through learned functions (*φ_e_, φ_h_, φ_x_*). Key operations include distance (dᵢⱼ)-based edge updates and equivariant coordinate transformations (highlighted).

We conducted extensive evaluations to demonstrate the effectiveness of YuelDesign across multiple dimensions. First, we assessed the chemical properties of the generated molecules, showing high drug likeness and low synthetic complexity. Second, we analyzed the chemical diversity of generated molecules, showing that YuelDesign explores novel chemical space while maintaining structural similarity to known ligands. Notably, the generated set included a broader range of functional groups, such as epoxides and nitriles, which were absent in native ligands in the training set. Third, we redocked the generated molecules to their target protein to evaluate protein-ligand interactions, and we found YuelDesign produces molecules with comparable docking energies to native ligands, especially for molecules with small size. Finally, we performed detailed analyses of the denoising process to understand how the model systematically refines molecular structures through atom type transitions, bond dynamics, and conformational changes.

## RESULTS

### Chemical Property Analysis of Diffusion-Generated Molecules

We conducted a comprehensive evaluation of the chemical properties of generated molecules, comparing YuelDesign with DiffSBDD and native ligands across multiple metrics (Figure 2, Table S1, Table S2). The analysis includes Quantitative Estimate of Drug-likeness^29^ (QED), Lipinski’s Rule of Five (RO5) compliance, Synthetic Accessibility Score^30^ (SAS), large ring formation rate, chemical validity, and connectivity.

**Figure 2.**
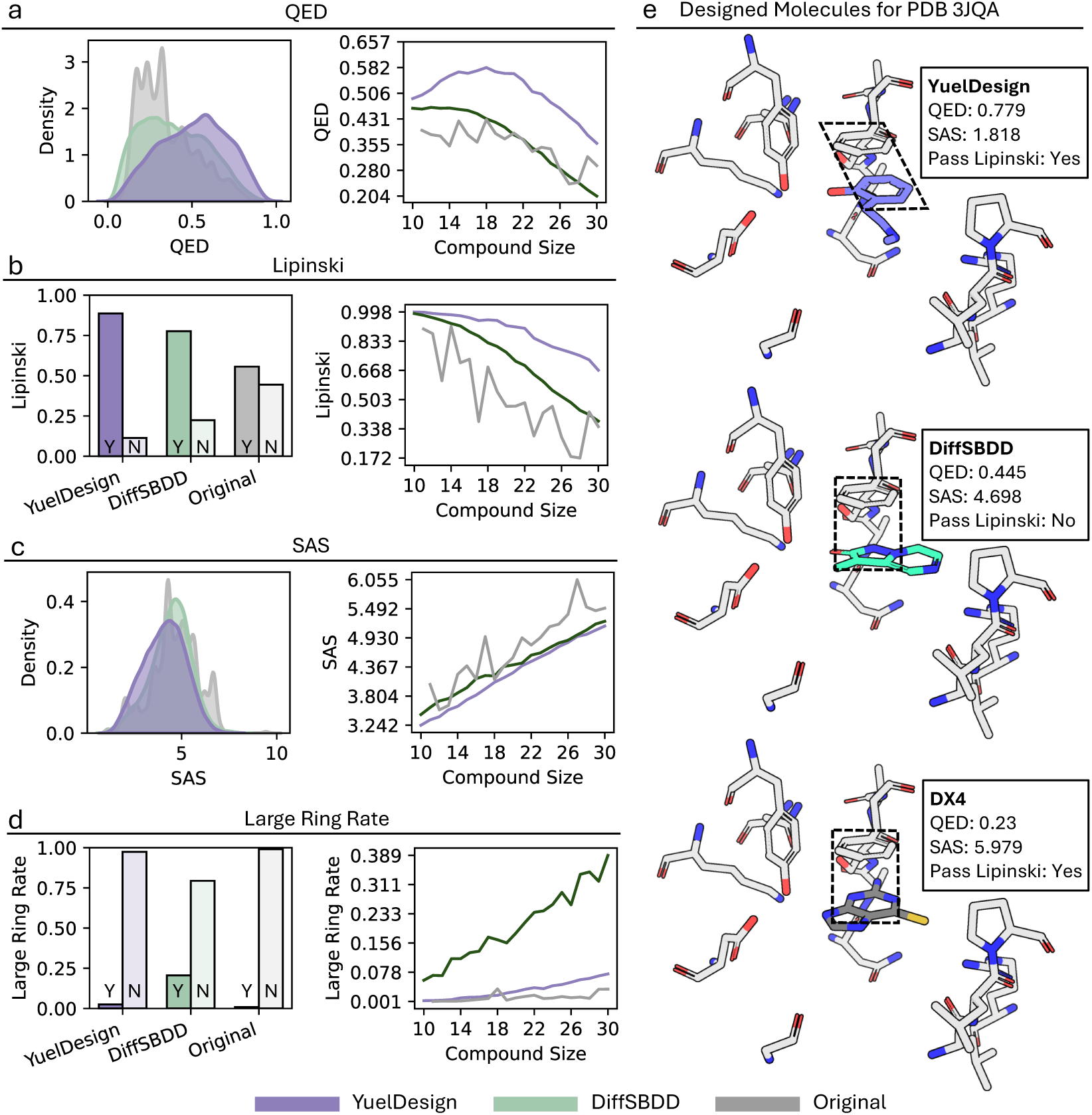
Evaluation of generative molecules and native ligands through multiple metrics. (a-d) Overall distribution of Quantitative Estimate of Drug-likeness (QED), Lipinski’s RO5 compliance (pass/fail ratios), Synthetic Accessibility Score (SAS), and Macrocycle (more than 6 atoms) formation rate, as well as their size dependence. (e) Comparison of molecules designed by YuelDesign, DiffSBDD, and the native ligands for Pteridine reductase 1 (PDB ID: 3JQA). Inside the dashed black box is the π–π stacking interaction. DX4 (2-amino-1,9-dihydro-6H-purine-6-thione) is the native ligand of 3JQA.

We evaluated the drug-likeness of generated molecules using two widely adopted metrics. QED is a composite score between 0 and 1 that combines multiple molecular descriptors, while Lipinski’s RO5 consists of four physicochemical parameter thresholds for bioavailability. YuelDesign-generated molecules consistently outperformed both DiffSBDD outputs and native ligands in terms of QED scores (Figure 2a, Table S3). QED values initially rose and then declined with molecular size, remaining higher than those of DiffSBDD. We also assessed RO5 compliance, and YuelDesign-generated molecules showed higher RO5 compliance rates than those produced by DiffSBDD and native ligands (Figure 2b, Table S4). RO5 compliance decreased with increasing molecular size.

We also evaluated the synthetic accessibility of generated molecules by using SAS, which ranges from 1 (easy to synthesize) to 10 (very difficult to synthesize). The SAS metric considers factors such as molecular complexity, presence of unusual structural features, and similarity to previously synthesized compounds. Synthetic accessibility analysis revealed that YuelDesign-generated molecules exhibited lower SAS values than both DiffSBDD outputs and native ligands (Figure 2c, Table S5). SAS values showed a positive correlation with molecular size.

Large rings, defined as cyclic structures containing more than 6 atoms, can pose significant challenges in molecular synthesis and affect drug-like properties. Only a small fraction of YuelDesign-generated molecules (2.5%) contained large rings, markedly lower than the 20.6% observed in DiffSBDD outputs (Figure 2d, Table S2). Even as molecular size increased, the proportion of large-ring structures in YuelDesign molecules remained below 10% (Table S6), demonstrating effective control over this undesirable feature.

We also compared validity (Figure S1) and connectivity (Figure S2, Table S1). Molecular validity serves as a fundamental measure of chemical feasibility, ensuring that generated structures adhere to basic chemical rules and constraints. The molecules generated with YuelDesign exhibited high validity rates, with 53.7% overall validity (Figure S1). As the size of molecule increases, the validity decreases. For molecules with only 10 heavy atoms, validity reaches 77.8%. Connectivity, which measures whether all atoms in a molecule are connected through a path of bonds, is a common challenge in diffusion-based molecular generation. This issue arises because diffusion models generate atomic coordinates independently, without explicit consideration of chemical bonding rules, leading to disconnected molecular fragments. The molecules generated with YuelDesign exhibited 99.5% overall connectivity (Figure S2). Connectivity decreases slightly with the increase of the size of generated molecules. Comparison with native ligands and DiffSBDD-generated molecules was not performed for validity and connectivity. DiffSBDD discards invalid and unconnected generated molecules inherently through post-generation filtering. Native ligands inherently exhibit 100% validity and connectivity.

Finally, we found that although side chains are not explicitly encoded as node features in the pocket’s graph representation, YuelDesign effectively captures interactions between generated compounds and protein side chains. For example, in the case of Pteridine reductase 1 (PDB ID: 3JQA), both YuelDesign and DiffSBDD generated molecules that exhibit π–π stacking interactions with the side chain of residue F97—mirroring the interaction observed in the native ligand (DX4) (Figure 2e). It implies that YuelDesign learns and restores critical protein–ligand interactions, even in the absence of explicit side chain information.

### Molecular Diversity Assessment of Diffusion Model Outputs

We conducted a thorough analysis of chemical diversity by examining the frequency distribution of functional groups in both generated molecules and native ligands (Figure 3a, Table S7). This analysis includes various structural motifs, such as aromatic rings, specialized ring systems, and functional groups that contain sulfur, nitrogen, and oxygen atoms. Additionally, we assessed the presence of common organic functional groups like acids, aldehydes, ketones, esters, amides, and other specialized moieties.

**Figure 3.**
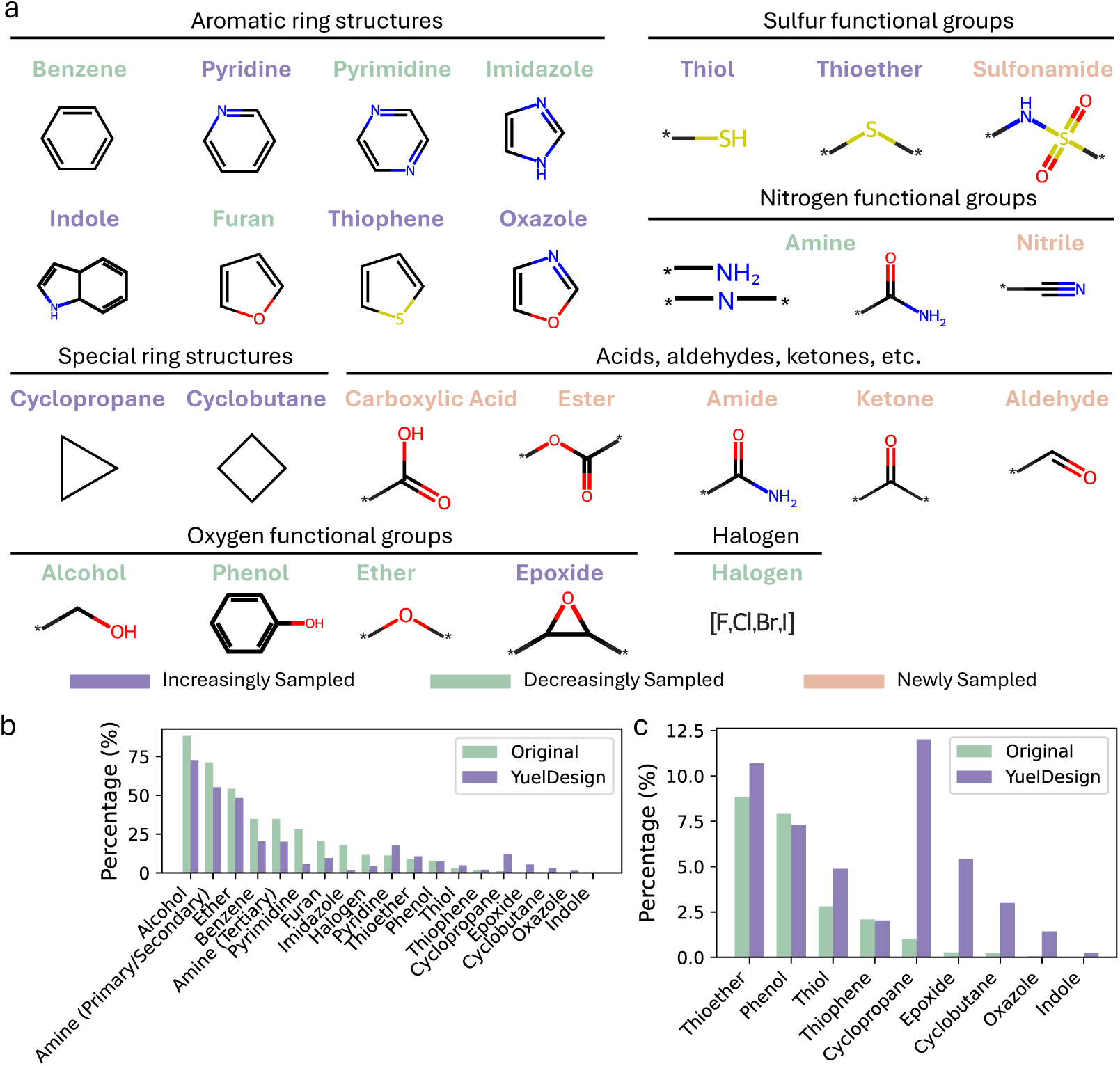
Functional group diversity analysis. (a) Representative structures of organic functional groups evaluated, categorized by chemical class (aromatic rings, N/S/O-containing groups, halogens, etc.). (b-c) Bar plots comparing the percentage distribution of functional groups in native ligands (green) versus YuelDesign (blue).

**Figure 4.**
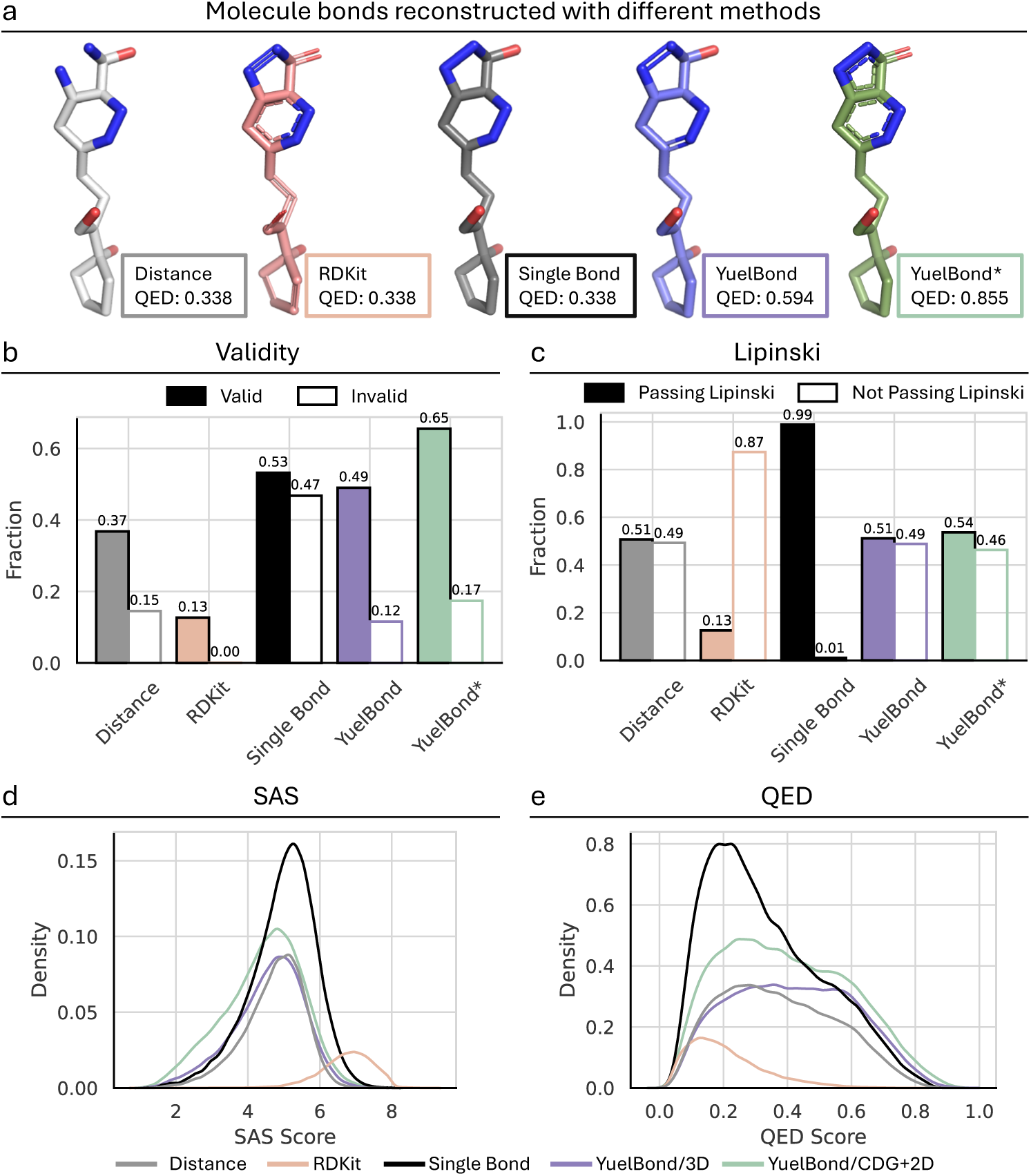
Comparative evaluation of bond reconstruction methods for small molecule generation. (a) Reconstructed molecular structures with associated QED scores (0-1 scale) for 5 methods: Distance-based, RDKit, Single Bond (baseline), YuelBond, and YuelBond*. (b) Fraction of valid versus invalid compounds for all 5 methods. (c) Fraction of molecules passing and not passing Lipinski’s Rule compliance. (d) Distribution of SAS for all 5 methods. (e) Distribution of QED for all 5 methods.

The analysis revealed distinct patterns in the distribution of functional groups between native ligands and generated molecules. Alcohol groups emerged as the most prevalent functional group in both sets, occurring in approximately 90% of native ligands and 70% of generated molecules. Amine groups ranked second in frequency, appearing in approximately 70% of native ligands and 55% of generated molecules. In contrast, several functional groups showed minimal representation, including epoxides, cyclobutanes, oxazoles, and indoles.

We observed a notable trend in the distribution patterns. Functional groups that were highly prevalent in native ligands (such as alcohols, amines, and ethers) exhibited reduced frequency in generated molecules. Conversely, uncommon functional groups in native ligands (including epoxides, cyclobutanes, oxazoles, and indoles) were sampled more frequently in the generated set. This shift is particularly evident in the case of epoxides, which increased from 0.1% in native ligands to 5.0% in generated molecules, suggesting that the model is exploring a broader chemical space beyond the constraints of the training data.

The analysis identified several new functional groups in the generated molecules, such as carboxylic acids, esters, amides, ketones, aldehydes, nitriles, and sulfonamides, that were not present in the native ligands in our training set. This increased chemical diversity highlights YuelDesign’s capability to create novel molecular structures while preserving chemical validity.

### Impact of Bond Reconstruction on Molecular Properties

Since the generated molecules may not have atom positions that lead to ideal bond lengths and angles, the bonds could be incorrect. We evaluated the effectiveness of various bond reconstruction methods on the generated molecules. Four distinct approaches were compared: (1) a baseline method using single bonds for all connections, (2) RDKit^31^, (3) a custom distance-based reconstruction algorithm (Methods), and (4) YuelBond^28^, a multimodal graph neural network framework specifically designed for bond order prediction in generated molecules. YuelBond operates in three modes: 3D (bond generation from precise 3D coordinates), CDG (reconstruction from <u>c</u>rude *<u>d</u>e novo* <u>g</u>enerated compounds), and 2D (reconstruction from all single bonds). Our primary analysis utilized the CDG mode, with additional testing of a combined approach (YuelBond*) that integrates CDG with the 2D mode.

We assessed the chemical properties of reconstructed molecules across metrics including validity, QED, Lipinski’s RO5 compliance, and SAS. YuelBond* demonstrated superior performance in validity, achieving a 65% success rate, while RDKit showed the lowest validity at 13%. This performance gap can be attributed to the fundamental differences in their design approaches. RDKit employs the xyz2mol^32^ program, which is widely used in chemical structure reconstruction but requires ideal bond lengths and angles. However, bond lengths and angles are often distorted in generated molecules. Similarly, the custom distance-based method faces the same limitation, which explains its lower validity rates.

Molecules reconstructed with single bonds demonstrated the highest Lipinski compliance rate (99%), followed by YuelBond* (54%), while RDKit achieved only 13% compliance. The superior performance of single-bond reconstruction aligns with Lipinski’s RO5, as single bonds tend to create molecules with fewer hydrogen bond donors and acceptors, and are lipophilic. The SAS and QED analyses show that YuelBond* resulted in lowest SAS and highest QED than the other methods. It suggests that while single-bond reconstruction maximizes Lipinski compliance rate, YuelBond* achieves a balanced trade-off by generating structurally diverse yet synthetically accessible and drug-like molecules.

### Protein-Ligand Interaction Analysis of Generated Molecules

We next conducted a comprehensive analysis of the protein-ligand interactions between generated molecules and their target proteins, comparing them with native ligands across multiple metrics (Figure 5), including binding affinity, structural stability, and molecular similarity assessments.

**Figure 5.**
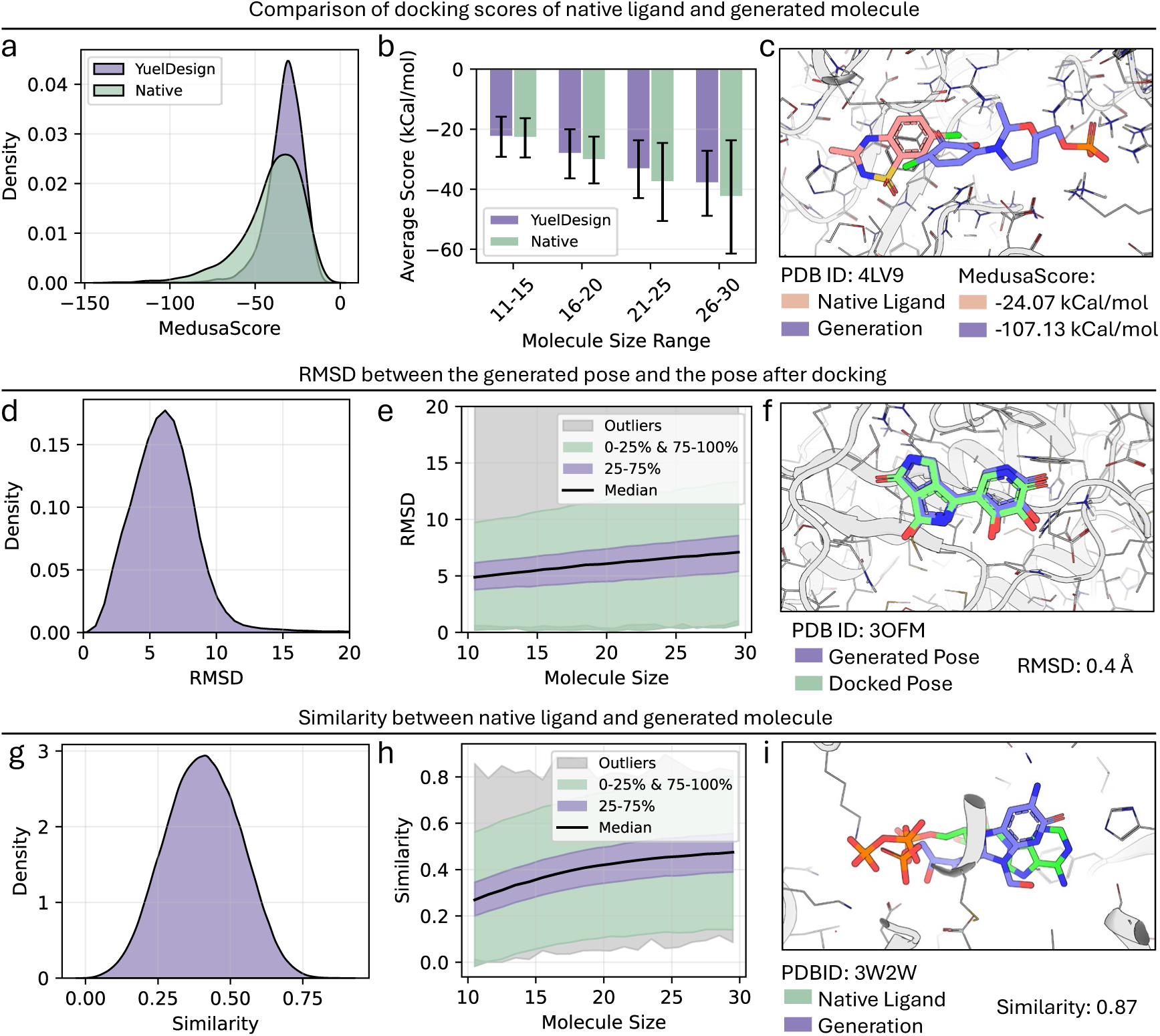
Comparative analysis of docking performance between generated molecules and native ligands across multiple metrics. (a-b) Distribution (a) and size dependence of MedusaScore (kcal/mol) of YuelDesign-generated molecules (blue) versus native ligands (green). (c) Example docking poses for PDB ID 4LV9. (d-e) Distribution (d) and size dependence (e) of RMSD between generated and redocking poses. (f) Structural alignment for PDB 30FM (RMSD=0.4 Å). (g-i) Distribution and size dependence of similarity scores between generated compounds and native ligands. (i) Overlay of native (green) and generated (blue) ligands for PDB 3W2W.

We redocked all the generated molecules to their targets with MedusaDock^20,21^. The docking score, MedusaScore^26,33^, is a physical energy-based metric that evaluates binding affinity, with lower scores indicating stronger binding. Analysis of the score distributions revealed that native ligands generally achieved lower scores compared to generated molecules, indicating higher binding affinity of native ligands to their targets (Figure 5a). This highlights the inherent challenge of generating small molecules with high binding affinity. However, for molecules with lower atom counts, the scores of generated molecules were comparable to native ligands (Figure 5b), suggesting that diffusion models can effectively generate high-affinity molecules in this size range. Notably, YuelDesign still produced many molecules with more favorable (lower) energy values than native ligands. An example of this enhanced binding was observed in human nicotinamide phosphoribosyltransferase (PDB ID: 4LV9), where the generated molecule achieved a MedusaScore of-107.13 kcal/mol, significantly more favorable than the native ligand’s - 24.07 kcal/mol (Figure 5c).

We also evaluated the conformational changes of generated molecules by analyzing RMSD after redocking with MedusaDock, which provides a measure of the accuracy of direct 3D structure generation using diffusion models. The distribution of RMSD values (Figure 5d) after redocking showed an average distance of 5-6 Å. While the RMSD may not seem low, it is important to note that even redocking of native ligands may result in some RMSD deviation from their crystallographic poses, as docking algorithms have inherent uncertainty in pose prediction. Therefore, the observed RMSD values reflect both the accuracy of our diffusion model and the inherent limitations of docking methods. We observed that the RMSD values tended to increase with molecular size (Figure 5e), likely due to the greater conformational flexibility of larger molecules. While most generated structures showed moderate deviations upon redocking, there were some notable exceptions - a small subset of molecules maintained very low RMSD values approaching 0 Å. For example, a molecule generated for protein kinase CK2 subunits CK2alpha prime (PDB: 3OFM) achieved an RMSD of only 0.4 Å after redocking (Figure 5f).

We also analyzed the similarity between generated molecules and native ligands using MACCS fingerprints as a metric (Figure 5g). The similarity distribution revealed that while some generated molecules showed high similarity to native ligands, such as the Cmr2-Cmr3 Subcomplex (PDB ID: 3W2W) (Figure 5i), many exhibited low similarity scores. This finding demonstrates the diffusion model’s ability to generate novel molecular structures while also producing compounds that maintain key features of known active ligands. The challenge of reproducing native ligands is expected, given the vast chemical space of potential binders for any given target. This pattern was consistent across different molecular sizes (Figure 5h).

### Case Studies of Target-Specific Small Molecule Generation

To further test the capability of YuelDesign in generating target-specific molecules, we conducted case studies on two important proteins: the dopamine receptor (PDB ID: 7CKZ) and serotonin receptor (PDB ID: 7E2Y). The protein structure of the dopamine receptor features a well-defined binding pocket with the native dopamine ligand forming specific interactions with key residues including TRP321 and SER202 (Figure 6a,b). YuelDesign successfully generated novel molecules with distinct pharmacophore patterns, as exemplified by the top docking score molecule (Figure 6c). The docking score distribution of generated molecules showed a mean binding energy of-34.54 ± 10.27 kcal/mol, which is more favorable than the native dopamine’s score of-30.60 kcal/mol (Figure 6d). Pharmacophore-based clustering analysis (Methods) revealed that the generated molecules explore diverse chemical space, with the top-scoring molecule exhibiting a significantly different pharmacophore pattern compared to dopamine (Figure 6e).

**Figure 6.**
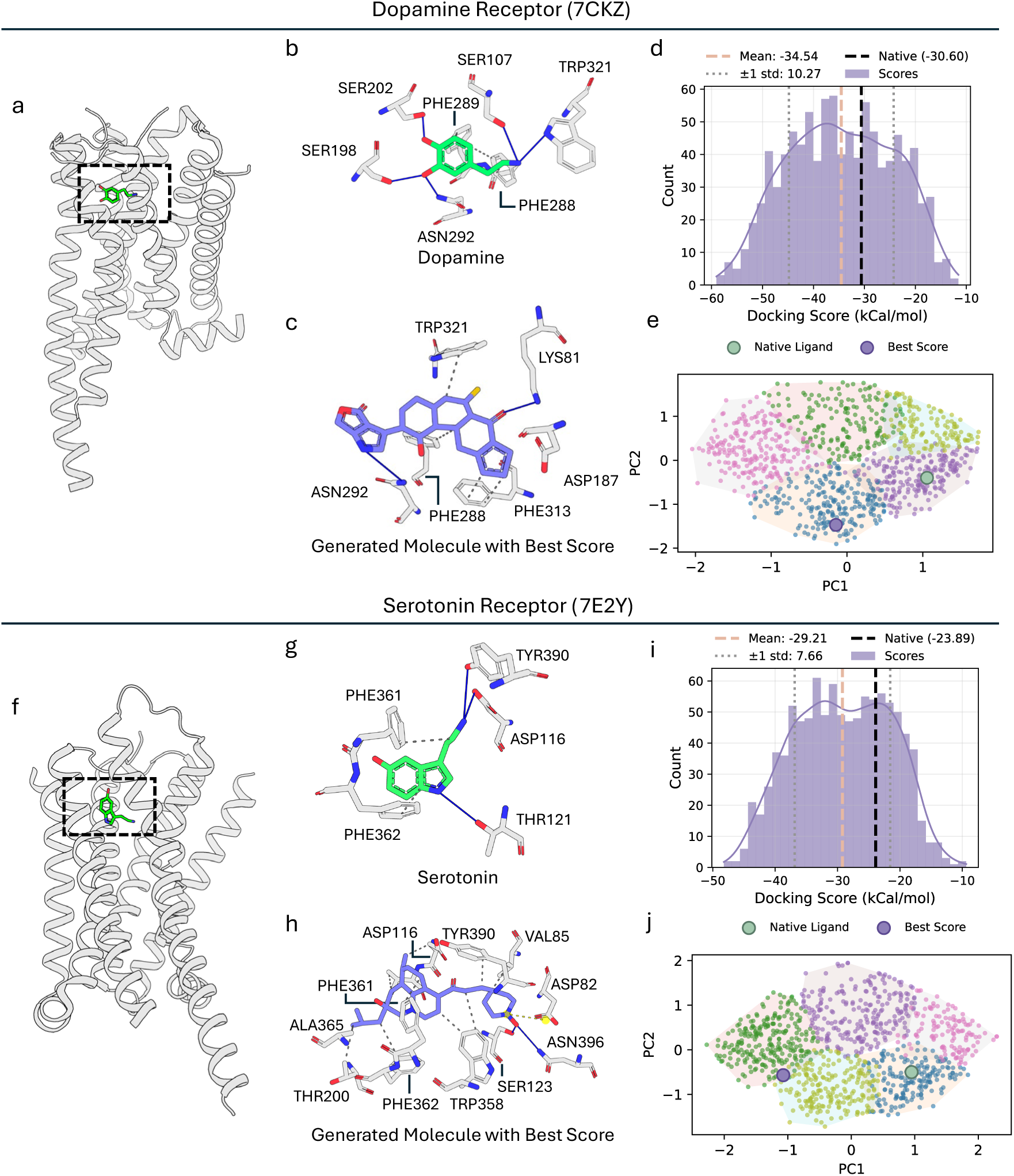
Case Studies of Target-Specific Small Molecule Generation. (a) Overall dopamine receptor (7CKZ) structure (gray cartoon) with native dopamine (green sticks). (b) Binding mode of dopamine (green) with residues (e.g., TRP321, SER202). (c) Top-scoring generated molecule (blue). (d) Docking score distribution (mean: −34.54 ± 10.27 kcal/mol; native: −30.60 kcal/mol). (e) Clustering of all generated molecules of 7CKZ according to their pharmacophore similarities. Different clusters are represented by different color regions. Dopamine is marked as green, and the best score molecule is marked as blue. (f) Serotonin receptor (7E2Y) structure binding with serotonine (green sticks). (g) Native serotonin binding pose. (h) Generated molecule. (i) Score distribution (mean: −29.21 ± 7.66 kcal/mol; native: −23.89 kcal/mol). (j) Clustering of all generated molecules of 7E2Y according to their pharmacophore similarities. Serotonin is marked as green, and the best score molecule is marked as blue.

The serotonin receptor (7E2Y) analysis yielded similar results (Figure 6f-j). The binding pocket accommodates the native serotonin ligand (Figure 6f,g), while YuelDesign generated novel molecules with distinct structural features (Figure 6h). The docking score distribution showed a mean binding energy of-29.21 ± 7.66 kcal/mol, comparable to the native serotonin’s score of-23.89 kcal/mol (Figure 6i). Pharmacophore clustering analysis demonstrated that the generated molecules explore novel chemical space, with the top-scoring molecule showing a different pharmacophore pattern compared to serotonin (Figure 6j).

### Structural Evolution During Denoising Process

Finally, we conducted a detailed analysis of the structural evolution during the denoising process of the diffusion model, examining both atomic and molecular-level changes (Figure 7). The analysis is for the methylmalonyl CoA mutase (PDB ID: 2REQ), and includes atom type transitions, structural stability, bond dynamics, and overall conformational changes.

**Figure 7.**
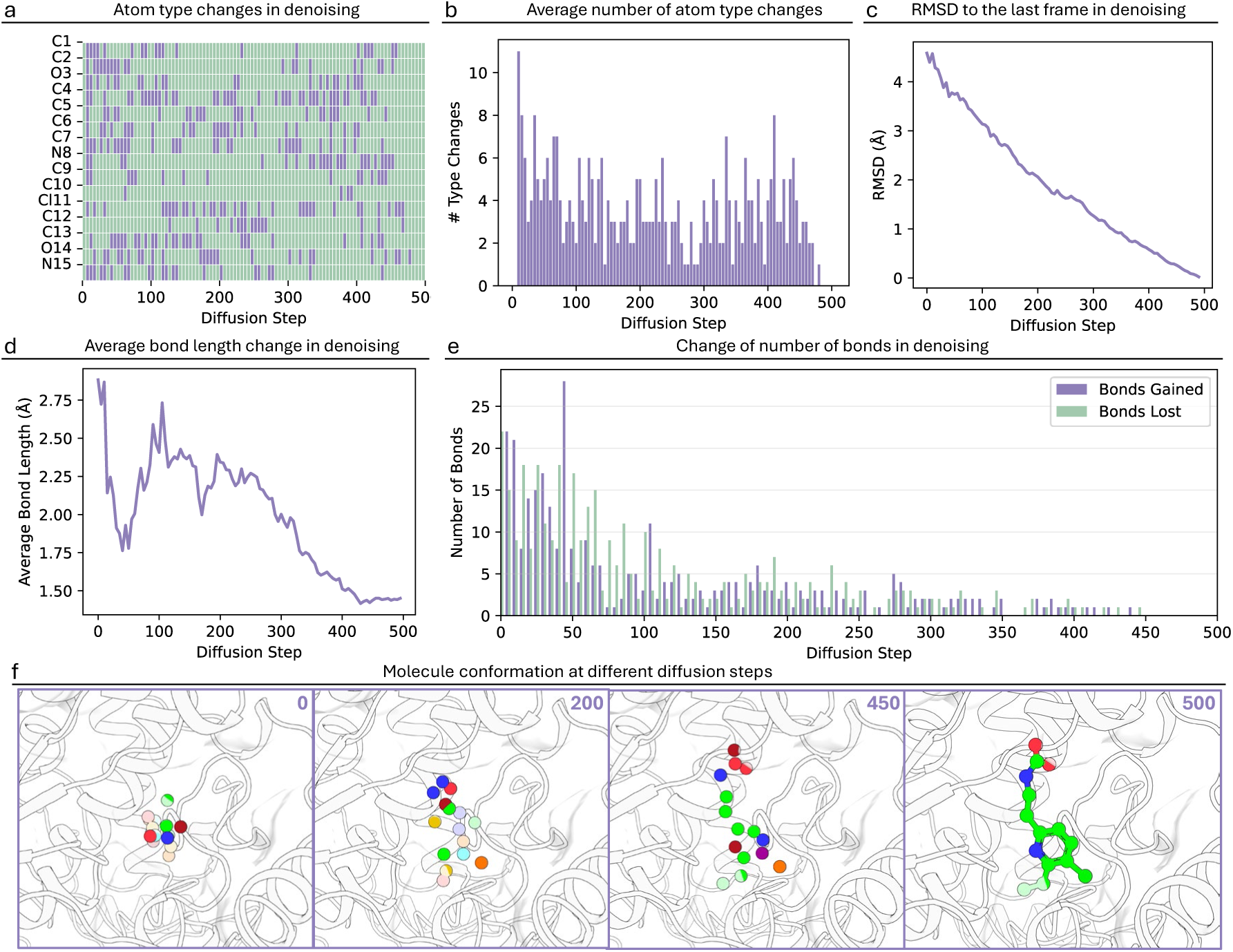
Structural evolution and conformational changes during the denoising process in the generation of PDB 2REQ. (a) Atom type changes across diffusion steps for individual atoms (C1-C15, O3, N6), where purple indicates atom type transitions and green represents stable atom types. (b) Average number of atom type changes per diffusion step. (c) Root Mean Square Deviation (RMSD) to the final frame during denoising. (d) Evolution of average bond length changes throughout the denoising process. (e) Quantification of bond formation (purple) and bond breaking (green) events at each diffusion step. (f) Visualization of molecular conformation at representative diffusion steps (0, 200, 450, and 500), showing the progressive refinement from initial random coordinates to the final chemically valid 3D structure. Colored spheres represent different atom types within the evolving.

Atom type evolution analysis demonstrated how atoms changed during the denoising process (Figure 7a). Individual atoms (C1-C15, O3, N6) showed varying degrees of stability, with some maintaining consistent atom types while others underwent transitions before reaching their final states. Notably, most atom types stabilized in the final 25 steps of diffusion, indicating that atom type determination primarily occurs in the later stages of the process. The average number of atom type changes per diffusion step showed a general decreasing trend (Figure 7b), with initial fluctuations that gradually diminished as the denoising process progressed toward the final molecular structure. This trend further confirms that atom types tend to stabilize in the final 25 steps of the diffusion process, suggesting a distinct phase where chemical identity becomes fixed.

We also calculated the RMSD between atomic coordinates at each denoising step and those in the final structure. Unlike atom type transitions, we found that RMSD showed a steady and uniform decrease throughout the process. We tracked the formation of bonds by analyzing bond lengths between atoms that form bonds in the final structure. Around step 200, these bond lengths began to consistently decrease, suggesting bond formation and subsequent optimization occurred at this stage. Additionally, we recalculated bonds at each diffusion step based on interatomic distances. In the early denoising phase, bond formation and breaking were highly frequent due to the close proximity of atoms sampled from Gaussian noise. Between steps 150-450, bond changes showed a gradually decreasing trend with fluctuations. In the final 50 steps, bond changes essentially ceased, indicating that the bonding pattern had stabilized.

The overall conformational evolution was visualized at key diffusion steps (0, 200, 450, and 500) (Figure 7f). The progression from initial random coordinates to the final chemically valid 3D structure was clearly evident. Notably, after step 450, the coordinates remained relatively stable, with subsequent steps primarily focused on fine-tuning atomic positions and adjusting atom types. This observation aligns with our earlier analysis of atom type transitions and bond dynamics, where we observed similar patterns of stabilization in the later stages of the diffusion process.

## DISCUSSION

The evaluation of YuelDesign shows several key findings about the capabilities and limitations of diffusion-based molecular design. The framework generates chemically valid and structurally stable molecules while maintaining properties relevant for drug discovery applications. The functional group distribution, particularly the balanced representation of common and rare groups, shows that YuelDesign can maintain similarity to known ligands while exploring novel chemical space. This balance is important for drug discovery, as it allows for the generation of both familiar and innovative molecular structures.

A key challenge in structure-based molecular design is handling protein pocket flexibility. Conventional approaches like DiffSBDD typically represent protein pockets using all atoms and construct graphs based on distance cutoffs. This approach has several limitations: first, due to the large number of atoms, these methods are forced to use distance-based cutoffs instead of fully connected graphs to maintain computational feasibility; second, the inclusion of side chain atoms may bias the model towards rigid conformations observed in crystal structures, limiting the exploration of novel binding modes. The use of distance cutoffs is particularly problematic in the context of flexible protein pockets, as it creates an artificial boundary for interactions that may change during the diffusion process. The graph structure remains fixed throughout the diffusion process, failing to capture the dynamic nature of protein-ligand interactions as atomic distances change during noising and denoising. For example, two atoms that are initially beyond the cutoff distance may come into proximity during the denoising process, but the model would not capture this interaction due to the fixed graph structure. This limitation is especially critical for protein-ligand interactions, where the binding process often involves significant conformational changes that bring distant atoms into contact.

To address these limitations, YuelDesign employs a strategy that focuses exclusively on the alpha carbon (CA) atoms of the protein backbone. To maintain the chemical identity of different amino acids, we encode each CA atom with a distinct type based on its amino acid identity, allowing the model to learn amino acid-specific interactions while still benefiting from the simplified backbone representation. This approach offers several advantages: first, it significantly reduces the total number of atoms in the system, making it computationally feasible to use fully connected graphs without distance cutoffs; second, it better represents the true nature of protein flexibility, as backbone movements are typically more constrained than side chain motions; and third, the fully connected graph architecture enables dynamic reweighting of all interatomic edges, providing a more accurate simulation of the evolving interaction landscape between proteins and ligands during denoising.

The bond reconstruction analysis shows the importance of specialized approaches for handling molecular connectivity. YuelBond’s performance compared to traditional methods like RDKit demonstrates the value of purpose-built solutions for generated molecules. The higher validity rates and better drug-likeness metrics achieved by YuelBond* suggest that combining different reconstruction modes can effectively balance chemical validity with property optimization. This finding has implications for the development of future molecular generation frameworks.

The protein-ligand interaction analysis provides insights into the binding properties of generated molecules. While native ligands generally exhibit lower energy values, YuelDesign can also generate molecules with comparable binding energies, particularly for smaller molecular sizes. The distribution of molecular similarity scores suggests that YuelDesign can explore the space between known ligands and novel structures, potentially leading to the discovery of new lead compounds.

Finally, the analysis of the denoising process shows how diffusion models refine molecular structures. The patterns in atom type transitions, bond dynamics, and conformational changes reveal a systematic approach to structure refinement. The stabilization of coordinates after step 450, combined with the continued fine-tuning of atomic positions and atom types, suggests that the model employs a hierarchical approach to structure generation. This understanding could guide future optimizations, such as decoupling atom type determination from 3D coordinate refinement, potentially improving denoising efficiency.

## METHODS

### Overview of the Molecular Generation Framework

YuelDesign employs a diffusion model for generating molecules. The model takes a protein pocket structure as input and generates molecules that fit within the pocket. The core of the model is built around an EGNN architecture that maintains rotational and translational equivariance, which is crucial for molecular design tasks. This equivariance ensures that the generated molecules maintain their physical properties regardless of their orientation in 3D space.

The workflow begins with data preparation, where the model processes the input pocket structure. The main model, implemented as a DDPM^34^ (Denoising Diffusion Probabilistic Model), consists of two key components: the EDM^35^ (Equivariant Diffusion Model) and the Dynamics module. The EDM handles the diffusion process, managing noise schedules and sampling, while the Dynamics module, built on EGNN, processes the molecular graph structure. During training, the model learns to denoise molecular structures by predicting the noise added to the molecule. The sampling process works in reverse, starting from random noise and gradually denoising to generate valid molecules.

After generating the 3D molecular structure, YuelBond is employed to reconstruct accurate chemical bonds. YuelBond analyzes the atomic coordinates and predicts bond orders using a multimodal graph neural network approach. It operates in CDG and 2D mode for *de novo* generated compounds, examining interatomic distances and chemical context to determine connectivity patterns. The final output is saved in both XYZ and PDB formats, with additional trajectory information if requested, allowing for detailed analysis of the generation process.

### Data Preparation

Binding MOAD^36^ (Mother of All Databases) is a carefully curated repository of high-quality protein–ligand crystal structures drawn from the Protein Data Bank^37^, designed to support structure-based drug discovery and polypharmacology investigations. The database has about 41,409 structural entries, with quantitative affinity measurements available for 15,223 (37%) of them, making it one of the most extensive resources for probing molecular recognition in biological systems. We prepared the MOAD dataset using a multi-step processing pipeline. For small molecules, we extract atomic features using RDKit, where each atom is represented by its 3D coordinates and a one-hot encoded feature vector. The atomic features include the atom type (C, N, O, F, S, Cl, Br, I, P) and its corresponding atomic number, which are encoded into a combined feature space. The protein pocket is processed using Biopython^38^, where we focus on the alpha carbon (CA) atoms of each residue to capture the essential structural information of the binding site. The use of CA atoms for protein representation is a deliberate choice to better model protein flexibility. By focusing on the backbone structure, we avoid the potential over-constraint that would come from including side chain atoms, which often adopt fixed conformations in crystal structures. Each CA atom is encoded with a distinct category based on its amino acid type, allowing the model to learn how different amino acids contribute to protein flexibility. This CA-only representation with amino acid-specific encoding allows the model to learn the essential backbone movements that characterize protein flexibility, while avoiding the bias towards rigid side chain conformations that might limit the exploration of novel binding modes.

The feature encoding scheme combines both molecular and protein information into a unified representation. For protein residues, we use a one-hot encoding scheme that includes 20 standard amino acids and several modified amino acids, cofactors, and nucleotides. The molecular atoms are encoded using a separate one-hot scheme for the allowed atom types. These two feature spaces are concatenated to create a unified feature representation, where the first part of the feature vector represents residue types, and the second part represents atom types. This combined feature space allows the model to distinguish between protein and molecular atoms while maintaining their distinct chemical properties.

The 3D structural information is preserved through the spatial coordinates of both protein and molecular atoms. For the protein pocket, we define the binding site as residues within 6 Å of any ligand atom, ensuring we capture the relevant protein-ligand interactions. The molecular and protein coordinates are concatenated to form a single coordinate matrix, while maintaining the distinction between pocket and molecular atoms through binary masks. This representation allows the model to learn both the local chemical environment and the global spatial arrangement of the binding complex.

The data is structured as a graph representation, where nodes represent atoms (both protein and molecular), and edges are defined for all node pairs. The fully connected graph structure, where every node is connected to every other node, is particularly important for modeling flexible protein pockets. This approach allows the model to capture long-range interactions and conformational changes in the protein structure, as atoms that may be distant in the initial structure but come into proximity during protein dynamics can still communicate through the graph edges. The edge features, which include spatial information and interaction types, enable the model to learn how protein flexibility affects molecular design.

To ensure computational efficiency and data quality, we implement several filtering criteria. Molecules with more than 150 total atoms (including both protein and molecular atoms) are excluded from the dataset. The data is processed in parallel using multiple workers, with each worker handling individual molecular entries independently. The final dataset is stored in a binary format, with each entry containing the complete molecular and pocket information, including spatial coordinates, feature vectors, and the necessary masks for graph construction. This structured format facilitates efficient data loading and processing during model training while preserving the essential 3D structural information of both the molecules and their binding pockets.

### Diffusion Model

The diffusion process utilizes a learned noise schedule that adapts to the complexity of the molecular design task. The model employs a combination of continuous and discrete features to represent molecular structures, where continuous features capture the 3D coordinates of atoms, and discrete features represent atom types and other molecular properties. The EGNN architecture processes these features through a series of message-passing layers, where each layer updates both the node features and their positions in 3D space while maintaining equivariance to rotations and translations.

During the generation process, the model starts with a random noise distribution and iteratively denoises it to produce a valid molecule. This process is guided by the pocket structure, which provides spatial constraints and interaction information. The training process involves optimizing the model to predict the noise added to molecular structures. The system’s output includes not only the final generated molecules but also the trajectory of the generation process, which can be valuable for understanding how the model arrives at its final structures.

The diffusion model in YuelDesign follows the principles of DDPM. The forward diffusion process gradually adds Gaussian noise to the molecular structure, while the reverse process learns to denoise and reconstruct the original structure.

The forward diffusion process is defined as:

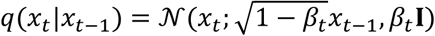

where *x_t_* represents the molecular structure at time step t, β_*t*_ is the noise schedule, and **I** is the identity matrix. The noise schedule follows a linear schedule:

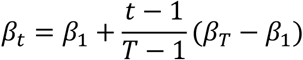

where T is the total number of diffusion steps, and β_1_ and β_*T*_ are the initial and final noise levels, respectively.

The reverse process is parameterized by a neural network that predicts the noise:

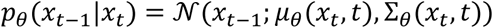

where *x_t−1_ and* μ_θ_(*x_t_, t*) are the mean and covariance predicted by the model. The training objective is to minimize the variational lower bound:

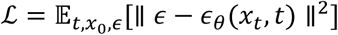

where ϵ is the added noise and ϵ_θ_ is the noise predicted by the model.

For molecular structures, we represent each atom by its 3D coordinates *r* ∈ ℝ^3^ and atom type ℎ ∈ ℝ^*d*^. The diffusion process is applied to both continuous (coordinates) and discrete (atom types) features:

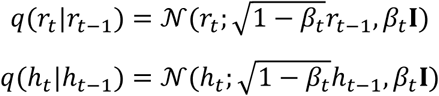

The model architecture employs EGNN to maintain rotational and translational equivariance:

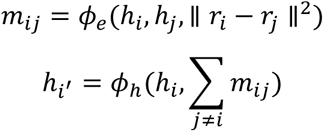

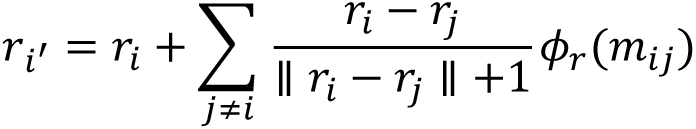

where ϕ_*e*_, ϕ_ℎ_, and ϕ_*r*_ are learnable functions implemented as neural networks.

### Implementation Details

The core architecture consists of 16 EGNN layers with a hidden dimension of 64, using SiLU activation functions. The model employs a fully connected graph structure, where every node in the molecular graph is connected to every other node, enabling comprehensive message passing between all atoms and capturing long-range interactions crucial for understanding molecular structure and properties. The model uses a learned noise schedule with L2 loss type and conditions on time, allowing the model to learn different behaviors at different stages of the diffusion process. The architecture includes data augmentation by randomly rotating the input structure to improve generalization. The model is trained with a learning rate of 2×10^-4^ and a batch size of 16, running for up to 1000 epochs. Performance is evaluated every 20 epochs, with 1 stability sample generated per evaluation. The model is implemented with Pytorch (version 2.6.0 with CUDA 12.4) and Pytorch Lightning (version 2.5.0).

### Bonds Reconstruction

We evaluated five distinct approaches for molecular bond reconstruction, each employing different strategies for determining molecular connectivity. The Single Bond method serves as our baseline approach, providing a simple reference point by replacing all bonds with single bonds regardless of atomic distances or chemical context. While chemically naive, this method offers valuable control for assessing the impact of more sophisticated bond assignment methods.

The Distance-based method employs a physics-informed approach that determines bond orders based on interatomic distances. This method utilizes predefined distance thresholds from bond length dictionaries, which contain typical bond lengths for different atom pairs. For example, for carbon-carbon bonds, the thresholds are 1.54 Å for single bonds, 1.34 Å for double bonds, and 1.20 Å for triple bonds; for carbon-nitrogen bonds, the thresholds are 1.47 Å (single), 1.29 Å (double), and 1.16 Å (triple); for carbon-oxygen bonds, the thresholds are 1.43 Å (single), 1.20 Å (double), and 1.13 Å (triple); and for nitrogen-nitrogen bonds, the thresholds are 1.45 Å (single), 1.25 Å (double), and 1.10 Å (triple). The method uses three margin parameters (0.10, 0.05, and 0.02 Å) to account for variations in bond lengths, with larger margins for single bonds (0.10 Å) and smaller margins for double (0.05 Å) and triple bonds (0.02 Å). By comparing actual distances between atoms against these thresholds, the method assigns appropriate bond orders (single, double, or triple).

RDKit represents a more sophisticated approach, leveraging RDKit’s built-in *DetermineBonds* functionality. This method combines distance-based rules with chemical knowledge to infer bond orders, incorporating additional chemical rules and heuristics to ensure chemically reasonable bond assignments. Its effectiveness is particularly notable in cases where atomic coordinates are well-defined and the molecular topology is relatively simple.

YuelBond stands as our primary bond prediction approach, based on a machine learning model trained to predict bond orders. YuelBond is a multimodal graph neural network framework specifically designed for bond order prediction in generated molecules. It addresses three key scenarios: (1) predicting bond orders from 3D coordinates, (2) reconstructing bonds from generated molecules with noisy geometries (crude *de novo* generated compounds, CDG), and (3) reassigning bond orders when initial molecular topologies are incorrect. YuelBond* represents combining the CDG and 2D mode of YuelBond.

### Chemical Properties Evaluation

The validity metric is a fundamental check that determines whether a molecule can be properly sanitized according to chemical rules. It is implemented using RDKit’s *SanitizeMol* function, which performs a series of chemical validity checks. A molecule is considered valid if it passes all these checks, which include proper atom valences, bond types, and overall chemical structure. The validity check is binary, returning True if the molecule is valid and False otherwise.

The connectivity metric evaluates whether all atoms in the molecule are connected through a path of bonds. This is implemented by constructing a graph representation of the molecule where atoms are nodes and bonds are edges. The metric uses NetworkX’s *is_connected* function to check if there exists a path between any pair of atoms in the graph. The connectivity check is also binary, returning True if the molecule is fully connected and False if there are disconnected components.

The large ring rate metric identifies molecules containing rings with more than 6 atoms. This is calculated using RDKit’s *GetSymmSSSR* (Smallest Set of Smallest Rings) function, which finds all rings in the molecule. The metric returns True if any ring has more than 6 atoms, and False otherwise. This is important because large rings are often undesirable in drug-like molecules due to their potential impact on molecular properties and synthetic accessibility.

The QED metric is a continuous measure that evaluates how drug-like a molecule is. It is calculated using RDKit’s *QED.default* function, which considers multiple molecular properties including molecular weight, logP, number of hydrogen bond donors and acceptors, and other structural features. The QED score ranges from 0 to 1, where higher values indicate more drug-like molecules. The formula for QED is:

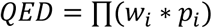

where *w_i_* are weights for different properties and *p_i_* are the property values normalized to [0,1].

The SAS (Synthetic Accessibility Score) metric estimates how difficult it would be to synthesize a molecule. It is calculated using a custom implementation that considers factors such as ring complexity, stereochemistry, and fragment contributions. The SAS score typically ranges from 1 to 10, where lower values indicate easier synthesis. The score is calculated as:

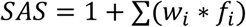

where *w_i_* are weights for different complexity factors and *f_i_* are the individual factor scores.

The Lipinski metric evaluates whether a molecule follows the RO5 criteria for drug-likeness. A molecule passes the Lipinski check if it satisfies all of the following conditions: molecular weight ≤ 500, logP ≤ 5, number of hydrogen bond donors ≤ 5, and number of hydrogen bond acceptors ≤ 10. The metric is binary, returning True if all criteria are met and False otherwise. This is implemented using RDKit’s *Descriptors* and *Lipinski* functions to calculate the individual properties.

### Functional Group Analysis

The functional group analysis employs a comprehensive set of SMARTS^39^ patterns to identify various chemical functional groups in the generated molecules. The analysis covers a wide range of functional groups including carboxylic acids, esters, amides, ketones, aldehydes, various types of amines, alcohols, phenols, ethers, epoxides, thiols, thioethers, sulfonamides, halogens, and various aromatic and special ring structures. Each functional group is defined using SMARTS, which are powerful molecular pattern-matching expressions. For example, carboxylic acids are identified using the pattern ‘C(=O)[OH]’, while primary/secondary amines use ‘[NX3;H2,H1;!$(NC=O)]’. The frequency calculation is performed by counting the occurrences of each functional group across all molecules in the dataset.

### Structural Analysis

The structural analysis of generated molecules includes three key aspects: bond length distributions, atom type transitions, and conformational changes. Bond length analysis focuses on the evolution of interatomic distances throughout the diffusion process. Using a 1.9 Å cutoff for covalent bond identification, the analysis tracks individual bond distances across all diffusion steps. Atom type transitions are analyzed through a systematic tracking of changes in atomic identities during the diffusion process. The analysis employs a binary matrix representation, where each element indicates whether an atom’s type changed at a particular diffusion step. The analysis also quantifies the frequency of type changes per step and identifies atoms that undergo the most transitions. Conformational analysis employs multiple metrics to assess structural evolution. The RMSD between intermediate and final structures quantifies the magnitude of conformational changes. Additionally, the analysis tracks bond dynamics by monitoring both the total number of bonds and the changes in bond counts between consecutive steps.

### Protein-Ligand Interactions Analysis

Docking energy is evaluated using MedusaDock scores, which quantify the strength of protein-ligand interactions. The analysis compares docking scores between generated molecules and native ligands across different molecular size ranges (e.g., 11-15, 16-20 atoms). The score distributions are visualized using kernel density estimation plots, where lower (more negative) scores indicate stronger binding affinity. This analysis helps understand how the model balances molecular size with binding strength. Molecular similarity analysis employs a multi-faceted approach using MACCS (Molecular ACCess System) fingerprints. These fingerprints encode molecular substructures as binary vectors, with similarity calculated using the Tanimoto coefficient to measure fingerprint bit overlap. The similarity analysis extends to examining how molecular size affects structural similarity. Data is organized into size-based bins and analyzed using box plots to show similarity score distributions. The results demonstrate that generated molecules maintain high similarity to target ligands across different size ranges, with median similarity scores typically exceeding 0.7. This analysis also identifies outliers and reveals the full range of similarity scores, providing insights into the model’s ability to balance structural conservation with innovation.

### Pharmacophore Clustering Analysis

To analyze the diversity and structural relationships among the generated molecules of the dopamine receptor and the serotonin receptor, we performed a pharmacophore-based clustering analysis. For each molecule, including both the generated candidates and the native ligand, we computed two-dimensional (2D) pharmacophore fingerprints using the RDKit cheminformatics toolkit. Specifically, we utilized the *Gobbi_Pharm2D* factory to generate pharmacophore fingerprints, which encode the presence and spatial arrangement of key pharmacophoric features such as hydrogen bond donors, acceptors, aromatic rings, and hydrophobic groups. Pairwise similarity between molecules was quantified by calculating the Tanimoto coefficient between their pharmacophore fingerprints. This resulted in a similarity matrix, where each entry represents the degree of pharmacophoric overlap between a pair of molecules. The similarity matrix was subsequently converted to a distance matrix (distance = 1 - similarity) to facilitate clustering. To group molecules with similar pharmacophoric profiles, we applied the KMeans^40^ clustering algorithm to the distance matrix. The number of clusters was set empirically as 5, and a unique random seed was used for each clustering run to ensure reproducibility. Each molecule was assigned to a cluster, allowing for the identification of major pharmacophoric chemotypes within the generated set. To visualize the clustering results, we performed principal component analysis^41^ (PCA) on the distance matrix, reducing the high-dimensional pharmacophore space to two principal components. The resulting coordinates were plotted, with each point representing a molecule and colored according to its assigned cluster. Convex hulls were drawn around each cluster to highlight their boundaries. The native ligand and the molecule with the best docking score were specifically highlighted to facilitate comparison with the generated candidates.

## ACKNOWLEDGMENTS

We acknowledge support from the National Institutes of Health 1R35 GM134864, the National Science Foundation grant 2210963, and the Passan Foundation. This project was supported by the Penn State College of Medicine’s Artificial Intelligence and Biomedical Informatics Program.

## DATA AND SOFTWARE AVAILABILITY

Source codes and test data are deposited at: https://bitbucket.org/dokhlab/yuel_design.

## SUPPORTING INFORMATION AVAILABLE

The supporting information provides Figure S1, Figure S2, and Tables S1-S7.

## AUTHOR CONTRIBUTIONS

Jian Wang contributed to the conceptualization, methodology, model development, data analysis, writing of the original draft, and visualization of the study. Dong Yan Zhang contributed to the writing of the original draft. Shreshty Budakoti contributed to the literature investigating. Nikolay V. Dokholyan provided supervision, resources, writing review and editing, project administration, and funding acquisition.

