## Supplemental Information for "A Diffusion-Based Framework for Designing Molecules in Flexible Protein Pockets"

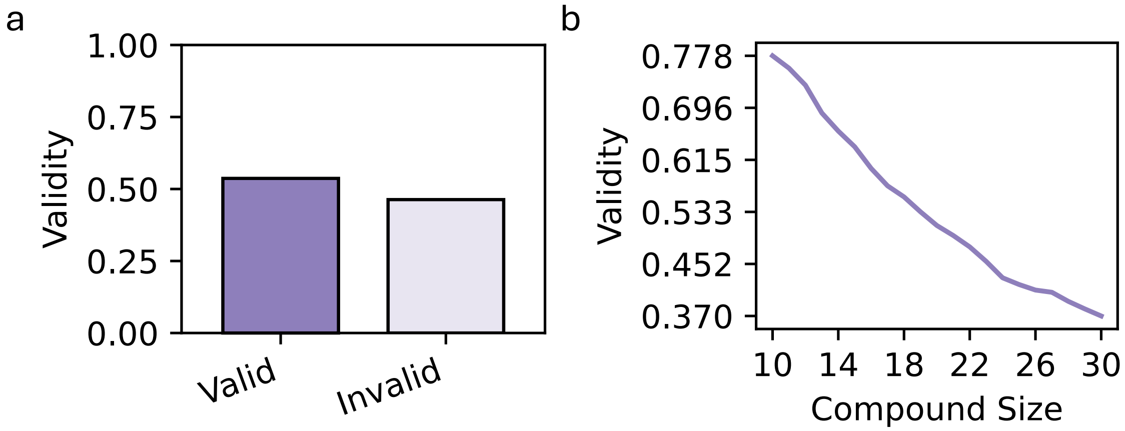


**Figure S1. The validity of molecules generated by YuelDesign.**

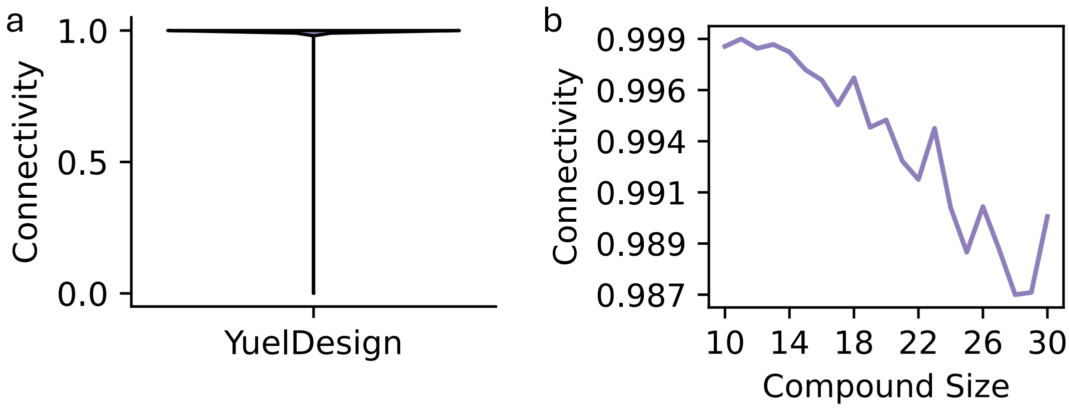


**Figure S2. The connectivity of molecules generated by YuelDesign.**

**Table S1. The validity and connectivity of molecules generated by YuelDesign.**

| Size | Validity | Connectivity |
|---|---|---|
| Overall | 0.537±0.499 | 0.995±0.073 |
| 10 | 0.778±0.416 | 0.998±0.039 |
| 11 | 0.758±0.428 | 0.999±0.034 |
| 12 | 0.732±0.443 | 0.998±0.040 |
| 13 | 0.688±0.463 | 0.999±0.038 |
| 14 | 0.660±0.474 | 0.998±0.043 |
| 15 | 0.635±0.481 | 0.997±0.052 |
| 16 | 0.601±0.490 | 0.997±0.056 |
| 17 | 0.574±0.494 | 0.996±0.066 |
| 18 | 0.557±0.497 | 0.997±0.055 |
| 19 | 0.533±0.499 | 0.995±0.074 |
| 20 | 0.512±0.500 | 0.995±0.071 |
| 21 | 0.496±0.500 | 0.993±0.084 |
| 22 | 0.478±0.500 | 0.992±0.089 |
| 23 | 0.456±0.498 | 0.995±0.074 |
| 24 | 0.430±0.495 | 0.991±0.096 |
| 25 | 0.420±0.493 | 0.989±0.106 |
| 26 | 0.411±0.492 | 0.991±0.096 |
| 27 | 0.407±0.491 | 0.989±0.106 |
| 28 | 0.393±0.488 | 0.987±0.115 |
| 29 | 0.381±0.486 | 0.987±0.115 |
| 30 | 0.370±0.483 | 0.990±0.098 |

Each value is represented as Mean±Standard Deviation.

**Table S2. Comparison of YuelDesign, DiffSBDD, and native ligands with respect to QED, SAS, Lipinski pass rate, and large ring formation rate.**

| | QED | SAS | Lipinski Pass Rate | Large Ring Rate |
|---|---|---|---|---|
| YuelDesign | 0.510±0.199 | 4.097±1.104 | 0.887±0.317 | 0.025±0.157 |
| DiffSBDD | 0.404±0.194 | 4.420±1.066 | 0.776±0.417 | 0.206±0.404 |
| Native Ligands | 0.351±0.165 | 4.645±1.234 | 0.556±0.497 | 0.009±0.094 |

Each value is represented as Mean±Standard Deviation.

**Table S3. Comparison of YuelDesign, DiffSBDD, and native ligands with respect to QED.**

| Molecule Size | YuelDesign | DiffSBDD | Native Ligands |
|---|---|---|---|
| 10 | 0.491±0.110 | 0.462±0.130 | N/A |
| 11 | 0.501±0.121 | 0.460±0.144 | 0.397±0.119 |
| 12 | 0.517±0.134 | 0.465±0.156 | 0.385±0.146 |
| 13 | 0.542±0.143 | 0.462±0.166 | 0.381±0.145 |
| 14 | 0.557±0.153 | 0.463±0.175 | 0.351±0.094 |
| 15 | 0.569±0.163 | 0.459±0.183 | 0.425±0.143 |
| 16 | 0.565±0.173 | 0.456±0.190 | 0.384±0.133 |
| 17 | 0.572±0.196 | 0.446±0.195 | 0.362±0.181 |
| 18 | 0.582±0.191 | 0.431±0.200 | 0.428±0.178 |
| 19 | 0.569±0.195 | 0.421±0.201 | 0.388±0.165 |
| 20 | 0.564±0.206 | 0.407±0.204 | 0.372±0.181 |
| 21 | 0.564±0.205 | 0.384±0.203 | 0.389±0.206 |
| 22 | 0.544±0.212 | 0.371±0.200 | 0.370±0.187 |
| 23 | 0.511±0.218 | 0.345±0.191 | 0.335±0.178 |
| 24 | 0.504±0.216 | 0.330±0.189 | 0.348±0.200 |
| 25 | 0.479±0.214 | 0.301±0.177 | 0.343±0.195 |
| 26 | 0.462±0.212 | 0.284±0.175 | 0.281±0.173 |
| 27 | 0.438±0.205 | 0.263±0.166 | 0.240±0.139 |
| 28 | 0.416±0.203 | 0.244±0.161 | 0.245±0.140 |
| 29 | 0.384±0.195 | 0.222±0.146 | 0.320±0.161 |
| 30 | 0.359±0.186 | 0.204±0.136 | 0.294±0.151 |

Each value is represented as Mean±Standard Deviation.

**Table S4. Comparison of YuelDesign, DiffSBDD, and native ligands with respect to Lipinski Rule of Five pass rate.**

| Molecule Size | YuelDesign | DiffSBDD | Native Ligands |
|---|---|---|---|
| 10 | 0.998±0.040 | 0.990±0.102 | N/A |
| 11 | 0.997±0.054 | 0.980±0.141 | 0.902±0.297 |
| 12 | 0.990±0.098 | 0.966±0.181 | 0.876±0.330 |
| 13 | 0.986±0.118 | 0.951±0.215 | 0.609±0.488 |
| 14 | 0.983±0.130 | 0.933±0.249 | 0.918±0.275 |
| 15 | 0.977±0.149 | 0.920±0.272 | 0.711±0.453 |
| 16 | 0.966±0.181 | 0.887±0.317 | 0.732±0.443 |
| 17 | 0.950±0.218 | 0.862±0.345 | 0.379±0.485 |
| 18 | 0.954±0.209 | 0.824±0.381 | 0.687±0.464 |
| 19 | 0.951±0.217 | 0.804±0.397 | 0.549±0.498 |
| 20 | 0.921±0.269 | 0.771±0.421 | 0.465±0.499 |
| 21 | 0.914±0.281 | 0.720±0.449 | 0.466±0.499 |
| 22 | 0.906±0.291 | 0.696±0.460 | 0.485±0.500 |
| 23 | 0.851±0.357 | 0.646±0.478 | 0.297±0.457 |
| 24 | 0.826±0.379 | 0.606±0.489 | 0.428±0.495 |
| 25 | 0.801±0.399 | 0.555±0.497 | 0.465±0.499 |
| 26 | 0.786±0.410 | 0.521±0.500 | 0.291±0.454 |
| 27 | 0.770±0.421 | 0.487±0.500 | 0.181±0.385 |
| 28 | 0.752±0.432 | 0.443±0.497 | 0.172±0.378 |
| 29 | 0.728±0.445 | 0.418±0.493 | 0.434±0.496 |
| 30 | 0.669±0.470 | 0.382±0.486 | 0.350±0.477 |

Each value is represented as Mean±Standard Deviation.

**Table S5. Comparison of YuelDesign, DiffSBDD, and native ligands with respect to SAS.**

| Molecule Size | YuelDesign | DiffSBDD | Native Ligands |
|---|---|---|---|
| 10 | 3.242±1.033 | 3.448±1.128 | N/A |
| 11 | 3.346±1.016 | 3.585±1.111 | 4.030±1.145 |
| 12 | 3.406±1.015 | 3.711±1.057 | 3.538±0.868 |
| 13 | 3.540±1.000 | 3.757±1.053 | 3.623±1.133 |
| 14 | 3.606±0.997 | 3.866±1.068 | 4.226±0.652 |
| 15 | 3.734±0.970 | 3.968±1.088 | 4.390±1.454 |
| 16 | 3.824±0.950 | 4.139±1.033 | 4.134±1.279 |
| 17 | 3.944±0.947 | 4.202±0.982 | 4.954±1.066 |
| 18 | 4.070±0.937 | 4.195±0.947 | 4.125±1.392 |
| 19 | 4.158±0.933 | 4.375±0.943 | 4.378±1.164 |
| 20 | 4.253±0.896 | 4.408±0.893 | 4.545±1.108 |
| 21 | 4.391±0.888 | 4.443±0.893 | 4.931±0.921 |
| 22 | 4.474±0.867 | 4.596±0.863 | 4.791±1.168 |
| 23 | 4.562±0.867 | 4.639±0.902 | 4.935±0.837 |
| 24 | 4.646±0.841 | 4.740±0.852 | 5.145±0.867 |
| 25 | 4.760±0.858 | 4.831±0.831 | 5.194±1.061 |
| 26 | 4.841±0.819 | 4.884±0.826 | 5.384±0.725 |
| 27 | 4.955±0.822 | 4.987±0.811 | 6.055±0.821 |
| 28 | 4.996±0.798 | 5.079±0.798 | 5.526±0.892 |
| 29 | 5.078±0.788 | 5.197±0.823 | 5.455±0.876 |
| 30 | 5.158±0.793 | 5.250±0.806 | 5.504±0.876 |

Each value is represented as Mean±Standard Deviation.

**Table S6. Comparison of YuelDesign, DiffSBDD, and native ligands with respect to large ring formation rate.**

| Molecule Size | YuelDesign | DiffSBDD | Native Ligands |
|---|---|---|---|
| 10 | 0.003±0.052 | 0.057±0.232 | N/A |
| 11 | 0.003±0.055 | 0.070±0.255 | 0.001±0.024 |
| 12 | 0.004±0.063 | 0.069±0.254 | 0.002±0.040 |
| 13 | 0.005±0.073 | 0.114±0.318 | 0.001±0.034 |
| 14 | 0.010±0.100 | 0.114±0.318 | 0.002±0.042 |
| 15 | 0.012±0.108 | 0.130±0.337 | 0.004±0.061 |
| 16 | 0.012±0.110 | 0.134±0.341 | 0.006±0.078 |
| 17 | 0.016±0.124 | 0.172±0.377 | 0.008±0.091 |
| 18 | 0.017±0.129 | 0.165±0.371 | 0.034±0.181 |
| 19 | 0.023±0.150 | 0.156±0.363 | 0.004±0.062 |
| 20 | 0.027±0.162 | 0.182±0.386 | 0.010±0.100 |
| 21 | 0.033±0.178 | 0.211±0.408 | 0.014±0.117 |
| 22 | 0.037±0.189 | 0.237±0.426 | 0.017±0.130 |
| 23 | 0.034±0.182 | 0.241±0.428 | 0.008±0.091 |
| 24 | 0.042±0.201 | 0.258±0.438 | 0.010±0.097 |
| 25 | 0.046±0.208 | 0.290±0.454 | 0.014±0.116 |
| 26 | 0.052±0.221 | 0.257±0.437 | 0.010±0.102 |
| 27 | 0.058±0.235 | 0.338±0.473 | 0.012±0.110 |
| 28 | 0.062±0.241 | 0.345±0.475 | 0.009±0.093 |
| 29 | 0.068±0.252 | 0.320±0.466 | 0.033±0.179 |
| 30 | 0.074±0.261 | 0.389±0.487 | 0.033±0.179 |

Each value is represented as Mean±Standard Deviation.

**Table S7. Chemical functional groups frequencies.**

| Functional Group | Percentage in Native Ligands | Percentage in YuelDesign-generated molecules | SMARTS Pattern | Description |
|---|---|---|---|---|
| Alcohol | 0.8831 | 0.7275 | [C&X4][O&H1] | Organic compound containing a hydroxyl group (-OH) attached to a carbon atom |
| Aldehyde | 0 | 0.0065 | [C&X3&H1](=O)C | Organic compound containing a carbonyl group (C=O) bonded to at least one hydrogen atom |
| Amide | 0 | 0.2316 | C(=O)N | Organic compound containing a carbonyl group (C=O) linked to a nitrogen atom, common in proteins and peptides |
| Amine (Primary/Secondary) | 0.7125 | 0.5526 | [N&X3;H2,H1;!$(NC=O)] | Organic compound containing nitrogen with one or two alkyl/aryl groups attached |
| Amine (Tertiary) | 0.3485 | 0.2013 | [N&X3]([#6])([#6])[#6] | Organic compound containing nitrogen with three alkyl/aryl groups attached |
| Benzene | 0.3487 | 0.2036 | C1CCCCC1 | Aromatic hydrocarbon with a six-membered ring containing alternating double bonds |
| Carboxylic Acid | 0 | 0.0072 | C(=O)[O&H1] | Organic compound containing a carboxyl group (-COOH), commonly found in amino acids and fatty acids |
| Cyclobutane | 0.0023 | 0.0299 | C1CCC1 | Cyclic hydrocarbon with a four-membered carbon ring |
| Cyclopropane | 0.0103 | 0.1202 | C1CC1 | Cyclic hydrocarbon with a three-membered carbon ring |

| | | | | |
|---|---|---|---|---|
| Epoxide | 0.0027 | 0.0543 | [C&R]1O[C&R]1 | Cyclic ether with a three-membered ring containing an oxygen atom |
| Ester | 0 | 0.1212 | C(=O)O* | Organic compound formed by the reaction of an acid with an alcohol, characterized by -COO- linkage |
| Ether | 0.5428 | 0.4823 | [O&D2]([#6])[#6] | Organic compound containing an oxygen atom connected to two alkyl or aryl groups |
| Furan | 0.2068 | 0.0952 | C1CCCO1 | Heterocyclic aromatic compound with an oxygen atom in a five-membered ring |
| Halogen | 0.116 | 0.0467 | [F,Cl,Br,I] | Element from group 17 (F, Cl, Br, I) that can form single bonds with carbon |
| Imidazole | 0.1789 | 0.0157 | C1CNC[N&H1]1 | Heterocyclic aromatic compound with two nitrogen atoms in a five-membered ring |
| Indole | 0.0001 | 0.0025 | C1CC2CCCC2[N&H1]1 | Heterocyclic aromatic compound containing a benzene ring fused to a pyrrole ring |
| Ketone | 0 | 0.4679 | C(=O)C | Organic compound containing a carbonyl group (C=O) bonded to two carbon atoms |
| Nitrile | 0 | 0.0205 | [C&!R]#N | Organic compound containing a cyano group (-C≡N) |
| Oxazole | 0.0007 | 0.0143 | C1COCNC1 | Heterocyclic aromatic compound containing both oxygen and nitrogen in a five-membered ring |
| Phenol | 0.0791 | 0.0729 | C1CCCCC1[O&H1] | Aromatic compound containing a hydroxyl group (-OH) directly attached to a benzene ring |

| | | | | |
|---|---|---|---|---|
| Pyridine | 0.1127 | 0.1776 | N1CCCCC1 | Heterocyclic aromatic compound with a nitrogen atom in a six-membered ring |
| Pyrimidine | 0.2833 | 0.0549 | C1CNCNC1 | Heterocyclic aromatic compound with two nitrogen atoms in a six-membered ring |
| Sulfonamide | 0 | 0.0061 | S(=O)(=O)N | Organic compound containing a sulfonyl group (-SO2-) linked to an amine |
| Thioether | 0.0884 | 0.107 | [#16]([#6])[#6] | Organic compound containing a sulfur atom connected to two alkyl or aryl groups |
| Thiol | 0.0281 | 0.0488 | [#16&H1] | Organic compound containing a sulfhydryl group (-SH) |
| Thiophene | 0.0209 | 0.0203 | C1CCSC1 | Heterocyclic aromatic compound with a sulfur atom in a five-membered ring |
